# PKR activation-induced mitochondrial dysfunction in HIV-transgenic mice with nephropathy

**DOI:** 10.1101/2022.10.03.510678

**Authors:** Teruhiko Yoshida, Khun Zaw Latt, Avi Z. Rosenberg, Briana A. Santo, Komuraiah Myakala, Yu Ishimoto, Yongmei Zhao, Shashi Shrivastav, Bryce A. Jones, Xiaoping Yang, Xiaoxin X. Wang, Vincent M. Tutino, Pinaki Sarder, Moshe Levi, Koji Okamoto, Cheryl A. Winkler, Jeffrey B. Kopp

## Abstract

HIV disease remains prevalent in the USA and chronic kidney disease remains a major cause of morbidity in HIV-1-positive patients. Host double-stranded RNA (dsRNA)-activated protein kinase (PKR) is a sensor for viral dsRNA, including HIV-1. We show that PKR inhibition by compound C16 ameliorates the HIV-associated nephropathy (HIVAN) kidney phenotype in the Tg26 transgenic mouse model, with reversal of mitochondrial dysfunction. Combined analysis of single-nucleus RNA-seq and bulk RNA-seq data revealed that oxidative phosphorylation was one of the most downregulated pathways and identified signal transducer and activator of transcription (STAT3) as a potential mediating factor. We identified in Tg26 mice a novel proximal tubular cell cluster enriched in mitochondrial transcripts. Podocytes showed high levels of HIV-1 gene expression and dysregulation of cytoskeleton-related genes; and these cells dedifferentiated. In injured proximal tubules, cell-cell interaction analysis indicated activation of the profibrogenic PKR-STAT3-platelet derived growth factor (PDGF)-D pathway. These findings suggest that PKR inhibition and mitochondrial rescue are potential novel therapeutic approaches for HIVAN.

**Translational Statement:** This work identified mitochondrial dysfunction in transgenic mice manifesting HIV-associated nephropathy mice kidney, using combination of single-nuclear and bulk RNA-seq analysis. Kidney damage was ameliorated by the PKR inhibitor C16, and mitochondrial rescue was shown by transcriptomic profiling and functional assay. These findings suggest that PKR inhibition and mitochondrial rescue are potential therapeutic approaches for HIV-associated nephropathy.

## Introduction

The World Health Organization (WHO) reported that in 2020, ∼1.5 million people became newly HIV positive, ∼37 million people were living with HIV, and ∼680,000 people globally died from HIV-related causes.^1^ HIV remains prevalent in the USA^2^ and even more highly prevalent in sub-Saharan Africa, especially in East and Southern Africa.^3^ Chronic kidney disease (CKD) remains a major co-morbidity in HIV positive individuals, even with availability of combined anti-retroviral therapy.^4^ The most severe form of CKD in persons with untreated or undertreated HIV infection is HIV-nephropathy (HIVAN), collapsing glomerulopathy. Therapy for HIVAN includes combined anti-retroviral therapy, coupled with renin-angiotensin-aldosterone system blockade, to prevent CKD progression.^5,6^

The Tg26 transgenic is a widely-used HIVAN model. The HIV-1 transgene expresses in kidney cells and the model manifests histological features similar to those of human HIVAN, including collapsing glomerulopathy, microtubular dilatation and interstitial fibrosis.^7^ The mice develop progressive renal dysfunction and progress to terminal uremia.^8,9^

Diverse therapeutic approaches are effective in Tg26 kidney disease, including inhibitors of mammalian target of rapamycin (mTOR),^10–12^ Notch inhibition,^13,14^ renin-angiotensin system inhibition,^15,16^ cyclin-dependent kinase inhibition,^17^ sirtuin1 agonist or overexpression,^18^ STAT3 activation reduction,^19,20^ retinoic acid receptor agonist,^21^ and NF-kB inhibition.^22^ Recent reports have identified several injury pathways, including NLRP3^23^ and mitochondrial dysfunction.^24,25^ Bulk multi-omics approaches using mRNA microarrays and protein-DNA arrays^26^ identified homeo-domain interacting protein kinase-2 (HIPK2) as a regulator of Tg26 renal pathology; this was confirmed in recent reports.^27,28^ A mRNA microarray approach characterized bulk transcriptional profiles during progressive renal disease.^29^ While these reports have provided novel insights into tissue transcriptional dynamics, studies have not been performed at single-cell or single-nucleus resolution.

Double-stranded RNA (dsRNA)-activated protein kinase (PKR) is a sensor for dsRNA and is activated in response to viral infections, including HIV-1. In the US and globally, HIV remains an important problem that disproportionately affects marginalized groups, including the Black/African-American community.^1^ *APOL1* risk variants, exclusively present in individuals with recent sub-Saharan ancestry, damage podocytes through various mechanisms including PKR activation by *APOL1* mRNA.^30^ The PKR-inhibiting oxoindole/imidazole compound C16 is beneficial in neuroinflammatory disease models.^31,32^

We hypothesized that PKR activation is a mechanistic pathway shared by HIV- and APOL1-mediated nephropathies, considering the high odds ratio for HIVAN among African Americans (OR 29) and South Africans (OR 89) carrying two *APOL1* risk alleles.^33^ We investigated the effects of PKR inhibition in the Tg26 HIVAN mouse model, which expresses HIV regulatory and accessory genes. We used single-nucleus and bulk RNA-sequencing methods to identify transcriptional changes between study groups and to uncover associated molecular mechanisms in an unbiased fashion.

## Methods

### Mice

Hemizygous (Tg26^+/-^) male mice were bred with wild type FVB/N female mice to generate Tg26 hemizygous mice. Transgenic mice were identified by PCR genotyping. We studied both male and female mice, aged 6–12 weeks. Mice in treatment groups were matched for sex.

For PKR inhibition treatment, the C16 treatment group mice received 10 µg/kg body weight of C16 (Sigma-Aldrich, St. Louis, MO) dissolved in 0.5% DMSO-PBS (10 ml/kg body weight), administered intraperitoneally three times weekly^30^ from 6 weeks to 12 weeks of age. Urine collection (in 24-hour metabolic cages) and body weight measurements were performed at 6 weeks of age before treatment and 12 weeks of age after treatment. Mice were euthanized and plasma, serum and kidney samples were collected at age of 12 weeks.

### Mouse kidney pathological evaluation, In situ hybridization (ISH) and Immunohistochemistry

Mouse kidney tissues were fixed with 10% buffered formalin for 24 hours, embedded in paraffin, and sectioned at 4-5 μm, and stained with hematoxylin and eosin, periodic acid Schiff, and picrosirius red. Chromogenic *in situ* detection of RNA was performed using RNAscope 2.5 HD Reagent Kit (catalog # 322310, Advanced Cell Diagnostics, Biotechne, Minneapolis, MN) with the RNA probes Mm-mt-Co1, Mm-mt-Atp6, dabB (negative control), Mm-Ppib (positive control) (catalog # 517121, 544401, 310043, 313911).

For immunohistochemistry, tissue sections were deparaffinized/rehydrated, antigens were retrieved by citrate buffer, and non-specific binding was blocked. Sections were incubated with primary antibody against phospho-Stat3 (Tyr705) (#9145, 1:100 dilution, Cell Signaling, Danvers, MA), phopho-PKR (Thr 446) (#sc16565, 1:50 dilution, Santa Cruz Biotechnology, Dallas, TX) and platelet-derived growth factor (PDGF)-D (ab181845, Abcam, 1:100 dilution, Cambridge, UK). Detailed methods including estimation of glomerular podocyte count by PodoCount^34^ are described in the Supplemental Methods.

### Immunoblotting

Tissues were lysed in a radioimmunopreciptation assay (RIPA) buffer (Thermo Fisher Scientific, Waltham, MA) containing a protease inhibitor/ phosphatase inhibitor cocktail (#78440, Thermo Fisher Scientific). Lysates were separated by SDS-polyacrylamide gel electrophoresis (SDS-PAGE) (gradient gel 4-12%, MOPS buffer) and the proteins subjected to Western blotting and blocked for one hour in Odyssey blocking buffer (LI-COR, Lincoln, NE). Blots were incubated following the iBind protocol (Thermo Fisher Scientific). Primary antibodies were Phospho-Stat3 (Tyr705) (#9145, 1:2000 dilution, Cell Signaling, Danvers, MA), Stat3 (#9139, 1:1000 dilution, Cell Signaling), β-actin (#47778, 1:5000 dilution, Santa Cruz Biotechnology), Total OXPHOS Rodent (#ab110413, 1:250 dilution, Abcam, Cambridge, UK), VDAC (# 4661, 1:1000 dilution, Cell Signaling). Blots were imaged using the Odyssey infrared scanner (LI-COR, Lincoln, NE).

### Bulk RNA-sequencing

Mouse kidney outer cortex tissues were dissected and homogenized in QIAzol (QIAGEN, Germantown, MD). Total RNA samples were extracted using RNeasy Plus Universal Kit (QIAGEN) following the manufacturer’s protocol including removal of genomic DNA step. RNA samples were pooled and sequenced on NovaSeq6000 S1 flowcell using Illumina TruSeq Stranded mRNA Library Prep and paired-end sequencing with read length 101bps (2×101 cycles). The samples had 46 to 72 million pass filter reads and more than 92% of bases calls were above a quality score of Q30. Sample reads were trimmed for adapters and low-quality bases using Cutadapt^35^. The trimmed reads were mapped to a reference genome (Mouse - mm10). Transcripts were annotated by Ensembl v96 using STAR aligner. Gene expression quantification analysis was performed for all samples using STAR/RSEM tools. DESeq2^36^ was used for differential expression analysis from raw count data and normalized data were used for Gene set enrichment analysis (GSEA v4.1.0).^37,38^ Pathway analysis including upstream regulator analyses were generated using QIAGEN Ingenuity Pathway Analysis.^39^

### Single-nucleus RNA-sequencing

Nuclei from frozen mouse kidney outer cortex tissue samples and glomeruli-enriched samples were prepared at 4°C.^40^ Tissue fragments (∼ 8 mm^3^) were cut by razor blades in EZlysis buffer (#NUC101-1KT, Sigma-Aldrich) and homogenized 30 times using a loose Dounce homogenizer and 5 times by tight pestle. After 5 min of incubation, homogenates were passed through 40 µm filters (PluriSelect, El Cajon, CA) and centrifuged at 500g at 4°C for 5 min. Pellets were washed with EZlysis buffer and again centrifuged at 500g at 4°C for 5 min. Pellets were resuspended in DPBS with 1% FBS and passed through 5 µm filters (PluriSelect) to make final nuclei preparations for loading on to 10xChromium Chip G (10x Genomics, Pleasanton, CA) and formation of gel beads in emulsion (GEM).

Single nuclear isolation, RNA capture, cDNA preparation, and library preparation were accomplished following the manufacturer’s protocol (Chromium Next GEM Single Cell 3’ Reagent Kit, v3.1 chemistry, 10x Genomics). Prepared cDNA libraries were sequenced. Analysis was performed with the Cell Ranger software using the default parameters with pre-mRNA analysis turned on. The reference was built from mm10 reference genome complemented with HIV-1 viral sequences.

### Single-nucleus RNA-seq Analysis

SoupX (version 1.5.2)^41^ was used to remove ambient RNA, following the default protocol by “autoEstCont” and “adjustCounts” functions. Doublets were identified and removed by DoubletFinder (version 2.0.3)^42^. Nuclei were filtered out that met any of the following criteria: detected genes < 200 or > 4000, total RNA count > 15000, or mitochondrial transcripts > 20%. Integration of single-nucleus gene expression data was performed using Seurat (version 4.0.5).^43^ After filtering, 57,061 cells remained. Clustering of the combined data used the first 30 principal components at a resolution of 0.6 and identified 25 distinct cell clusters. After removal of 2 doublet clusters, 56,976 cells from 23 clusters were used for downstream analysis. Cell type identification was done based on the expression levels of known marker genes. Pseudotime analysis of podocytes and proximal tubule cells was performed by using the R package Monocle 3 (version 1.0)^44^, considering wild-type cells as the root state. Cell-cell interaction analysis was performed using CellChat (version 1.5.0).^45^ Pathway analysis including upstream regulator analysis were accomplished through the use of QIAGEN Ingenuity Pathway Analysis.^39^

### Seahorse Extracellular Flux Assay

Seahorse 96-well assay plates (Agilent, Santa Clara, CA) were pre-coated twice with 20 μl/well of 0.01% poly-L-lysine solution (P4707, Sigma-Aldrich) and washed twice with PBS, 200 μl/well. Glomerular and proximal tubular samples were plated in EGM-2 medium (CC-3162, Lonza, Walkersville, MD) and placed in a CO_2_-incubator for 30 minutes for the attachment.

Seahorse Mito Stress Tests were conducted as described.^30^ Data were normalized by total nucleic acid content measured by CyQUANT Cell Proliferation (C7026, Thermo Fisher Scientific) and analyzed by Wave 2.6.1 (Agilent). Detailed method is described in the Supplemental Methods.

### Study Approval

Mouse experiments were conducted in accordance with the NIH Guide for the Care and Use of Laboratory Animals and were approved in advance by the NIDDK Animal Care and Use Committee (Animal study proposals, K097-KDB-17 & K096-KDB-20).

## Results

### PKR inhibition ameliorates kidney injury in Tg26 mice

The trans-activating regions (TAR) RNA in the HIV-1 long terminal repeats (LTRs) at the 5’ and 3’ ends of the HIV genome form double stranded RNA (dsRNA) structures and induce PKR autophosphorylation, thereby activating PKR.^46^ As PKR is a potent driver of many stress response pathways, including translational shutdown, apoptosis, inflammation and metabolism, we hypothesized that PKR inhibition by the PKR-specific oxoindole/imidazole inhibitor C16 might rescue kidney injury in Tg26 mice. We administered C16 to Tg26 and wild-type mice from 6 to 12 weeks of age, and we evaluated the kidney phenotype. C16-treated Tg26 mice had lower serum creatinine (**Figure 1A**), reduced albuminuria (**Figure 1B, 1C**) and reduced pPKR abundance (**Supplemental Figure 1A-D**) compared to control Tg26 mice. Further, C16 treatment reduced urinary excretion of the kidney injury marker neutrophil gelatinase-associated lipocalin (NGAL) (**Figure 1D**).

**Figure. 1.**
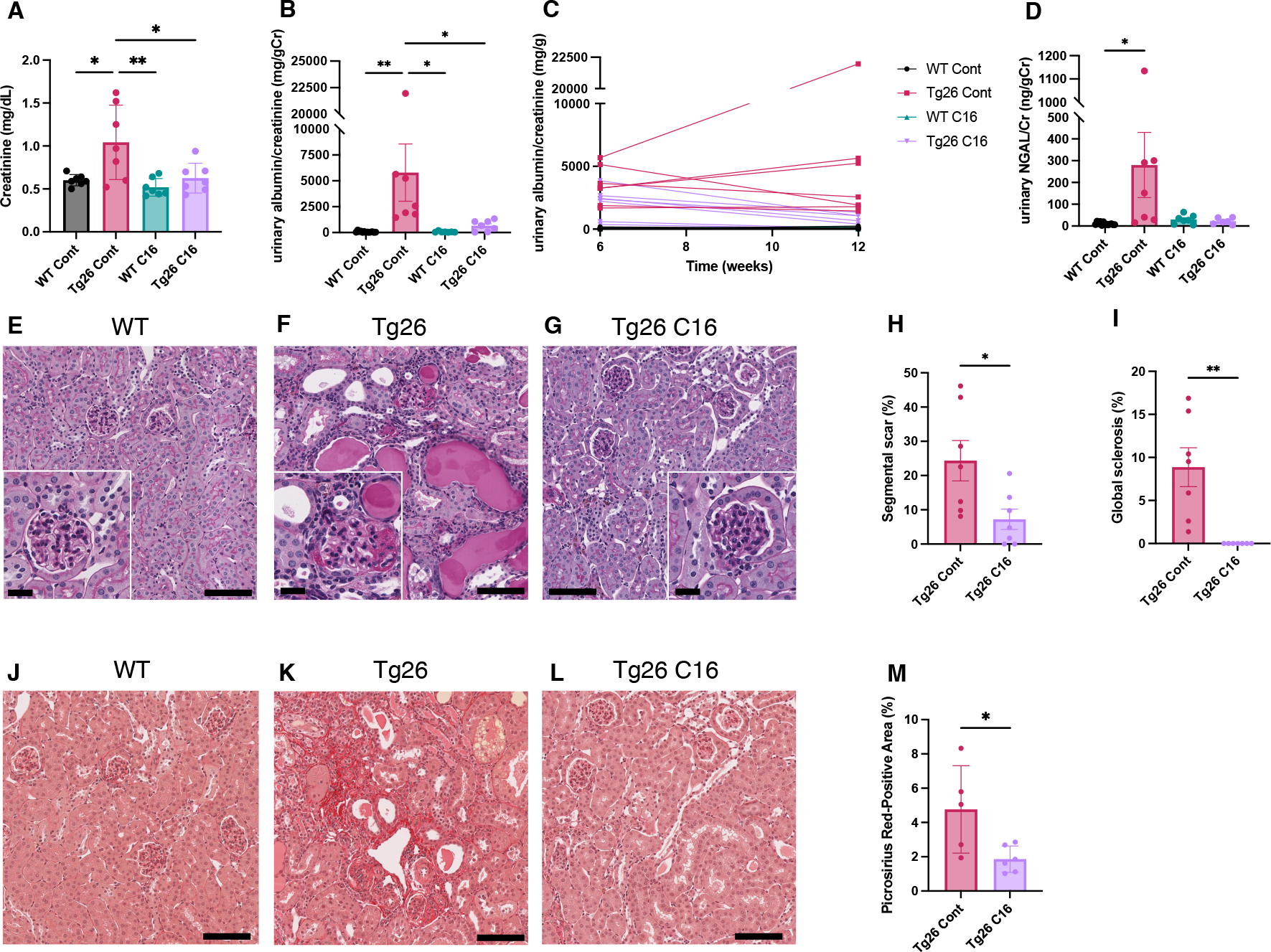
PKR inhibition by C16 ameliorates Tg26 mice kidney phenotype. (**A-D**) Shown are the following: plasma creatinine (mg/dL), urinary albumin-to-creatinine ratio (mg/g creatinine), urinary albumin-to-creatinine ratio (mg/g creatinine) of 6 and 12 weeks of age, urinary NGAL-to-creatinine ratio (ng/g creatinine). (**E-G**) Representative PAS staining images of WT, Tg26 and C16 treated Tg26 kidney. (**H, I**) Quantitative analysis of glomeruli for segmental scarring and global sclerosis. (**J-L**) Representative Picrosirius Red-staining images of WT, Tg26 and C16 treated Tg26 kidney. (**M**) Quantitative analysis of Picrosirius Red-staining area. (One-way ANOVA (**A, B, D**), t-test (**H, I, M**); *, P<0.05; **, P<0.01; scale bars are 50 μm)

At 12 weeks of age, there was less glomerulosclerosis and of microtubular dilatation in C16-treated Tg26 mice (**Figure 1E-G**). Histomorphologic quantification confirmed that C16 treatment reduced glomerular injury, assessed as segmental glomerulosclerosis and global glomerulosclerosis (**Figure 1H, 1I**), and as fibrosis extent, quantified by picrosirius red staining (**Figure 1J-M**).

### Combination of single-nucleus RNA-seq and bulk RNA-seq to profile transcriptomic changes in Tg26 mice

To investigate molecular mechanisms in Tg26 kidney and the effect of PKR inhibition, we conducted bulk RNA-seq of kidney cortex from the four groups (WT, WT treated with C16, Tg26, and Tg26 treated with C16; n=3 each) and single-nucleus RNA-seq of kidney cortex from six samples (WT, Tg26, and Tg26 treated with C16; n=2 each) and of two samples of glomeruli (WT and Tg26; n=1 each) to enrich for glomerular cells (**Figure 2A**). Bulk RNA-seq data clustered well by treatment groups in a principal component analysis plot (**Figure 2B**). Single-nucleus RNA-seq profiled 56,976 nuclei. We identified 23 cell clusters, including a novel cell type (PT-Mito, proximal tubule cell cluster with a higher expression level of mitochondrial genes compared to adjacent cells), as shown as a uniform manifold approximation and projection (UMAP) plot (**Figure 2C, Supplemental Figure 2C**). Marker genes for each cluster used for annotation are shown in **Figure 2D**. Taking advantage of unbiased clustering in single-nucleus RNA-seq, we tabulated cell numbers from each mouse kidney, using kidney cortex and glomeruli (**Figure 2E**, **Supplemental Figure 3A**). The gene encoding PKR, *Eif2ak2*, was expressed in glomeruli and all tubular segments (**Supplemental Figure 3B**).

**Figure. 2.**
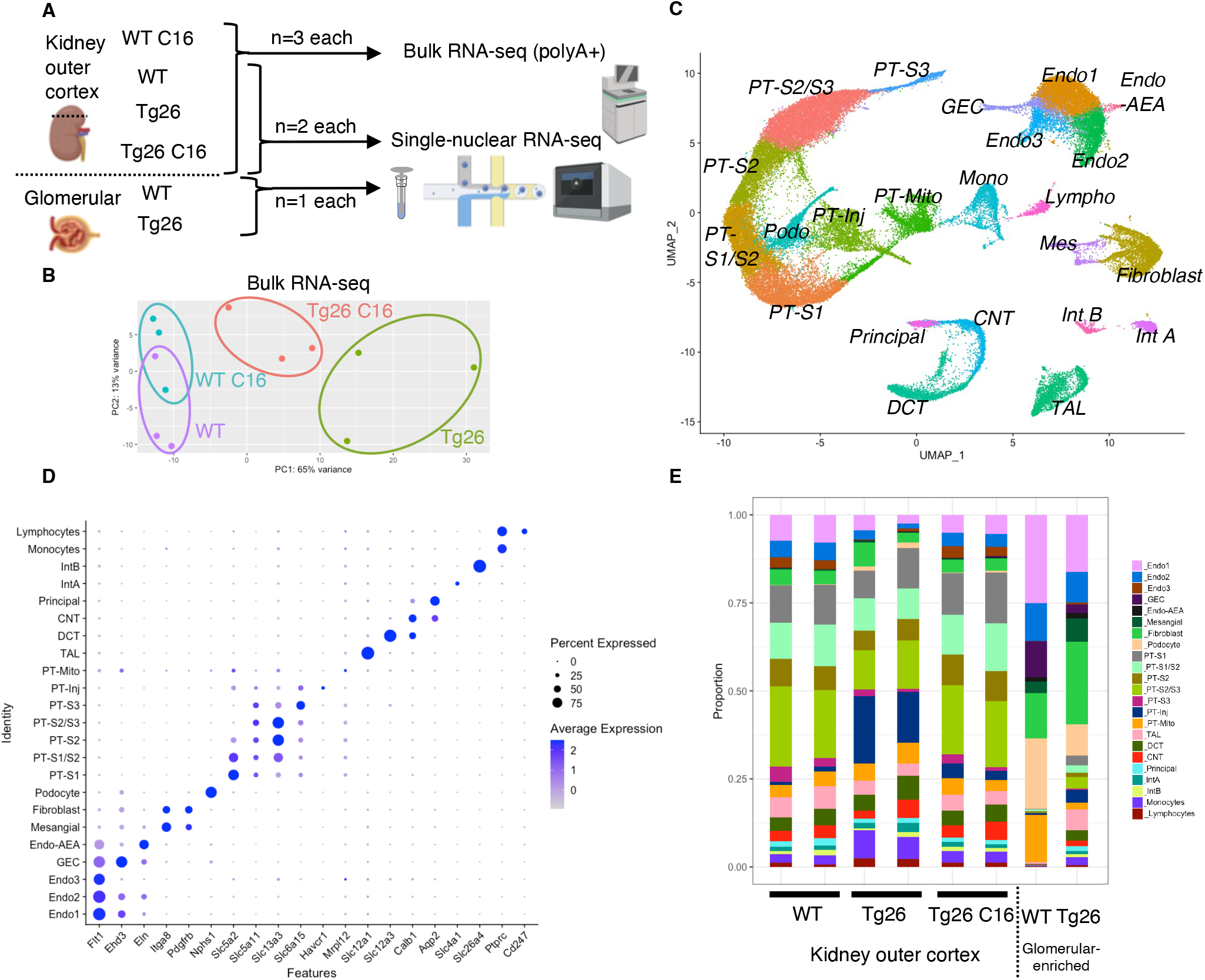
Overview of bulk RNA-seq and single-nucleus RNA-seq experiments. (**A**) Shown is the workflow of the bulk RNA-seq and single-nucleus RNA-seq experiments. (**B**) Principal component analysis plot of bulk RNA-seq results. (**C**) UMAP plot of single-nuclear RNA-seq data from 8 samples, 56,976 nuclei, showing 23 clusters. (**D**) Shown is a dot plot of 23 marker genes, each characteristic for the cluster. (**E**) Shown is the ratio of nuclei grouped to each cluster in each sample.

### Downregulation of mitochondrial genes in Tg26 kidneys

GSEA results of bulk RNA-seq showed that mitochondrial-related pathways were the most downregulated pathway in Tg26 mice when compared with WT mice, suggesting that mitochondrial gene transcription (for both nuclear-genome and mitochondrial genome-encoded genes) was significantly downregulated (**Figure 3A**). Expression levels of specific mitochondrial genes were assessed (**Figure 3B**) and showed that C16 treatment reversed the downregulation of these mitochondrial genes (**Figure 3B, 3C**). Western blot analyses were consistent with bulk RNA-seq results showing lower abundance of mitochondrial complex I and II in Tg26 kidney and this was restored to normal by C16 treatment (**Figure 3D, Supplemental Figure 4A-D**). Mitochondrial DNA copy numbers were decreased in Tg26 mice, but C16 treatment did not reverse these changes (**Supplemental Figure 4E, 4F**).

**Figure. 3.**
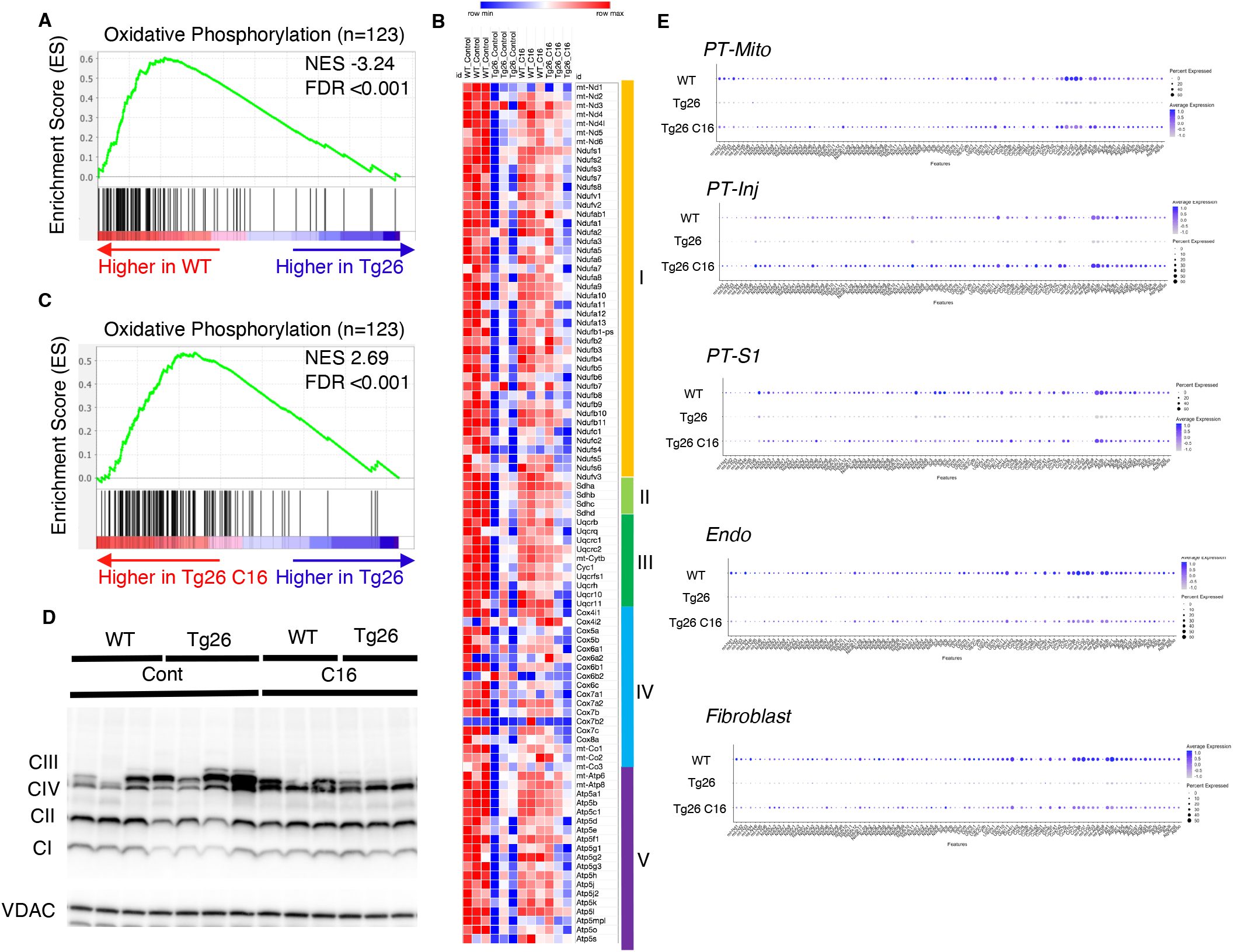
Oxidative phosphorylation genes are downregulated in Tg26 mice and downregulation is reversed by PKR inhibition using C16. (**A**) Shown is the enrichment plot of oxidative phosphorylation pathway based on bulk mRNA-seq comparing Tg26 and WT. (**B**) Shown is the enrichment plot of oxidative phosphorylation pathway based on bulk mRNA-seq comparing C16 treated Tg26 and Tg26. (**C**) Heatmap of expressed genes in oxidative phosphorylation pathway (n=123), based on data from bulk mRNA-seq. (**D**) Western blot to identify mitochondrial subunits CI through CIV and VDAC. (**E**) Dot plot showing expression of oxidative phosphorylation pathway genes in PT-Mito, PT-S1, PT-Inj, Endo, and Fibroblast cluster by snRNA-seq.

These findings suggest that PKR inhibition by C16 rescued transcriptional downregulation of both nuclear-encoded and mitochondrial-encoded mitochondrial genes, but the rescue was not through increased mitochondrial DNA copy number. Single-nucleus RNA-seq and subsequent pathway analysis showed that the majority of cell types, including mitochondrial proximal tubules (PT-Mito), PT-S1, injured PT (PT-Inj), and endothelial cells, manifested global downregulation of mitochondrial-expressed genes in Tg26 kidneys and that this decline was rescued by C16 treatment (**Figure 3E**).

### Novel proximal tubular cell cluster with high mitochondrial gene expression

Based on unbiased clustering, we identified a distinct proximal tubule cell cluster characterized by higher expression level of mitochondrial genes (PT-Mito) and consisting of 3.1-5.8% of kidney cortex cells (**Figure 2E**). We analyzed the seven proximal tubular cell clusters and found distinctive markers that distinguish each cluster **(Figure 4A**). Based on UMAP, this PT-Mito cluster was in proximity to proximal tubule-segment1 (PT-S1) and proximal tubule-segment2 (PT-S2) (**Figure 2C**). To confirm the presence of cells giving rise to this cluster, *in situ* hybridization (ISH) of mt-*Co1* and mt-*Atp6* were performed. We observed transcripts inside some nuclei in many putative PT-Mito segments with high expression of these genes (**Figure 4B, 4C, Supplemental Figure 5A, 5B**). We also confirmed the existence of similar PT-Mito cluster in published human kidney single-nuclear RNA-seq data^47^ by the re-analysis of the original data (**Supplemental Figure 6A-C**).

**Figure. 4.**
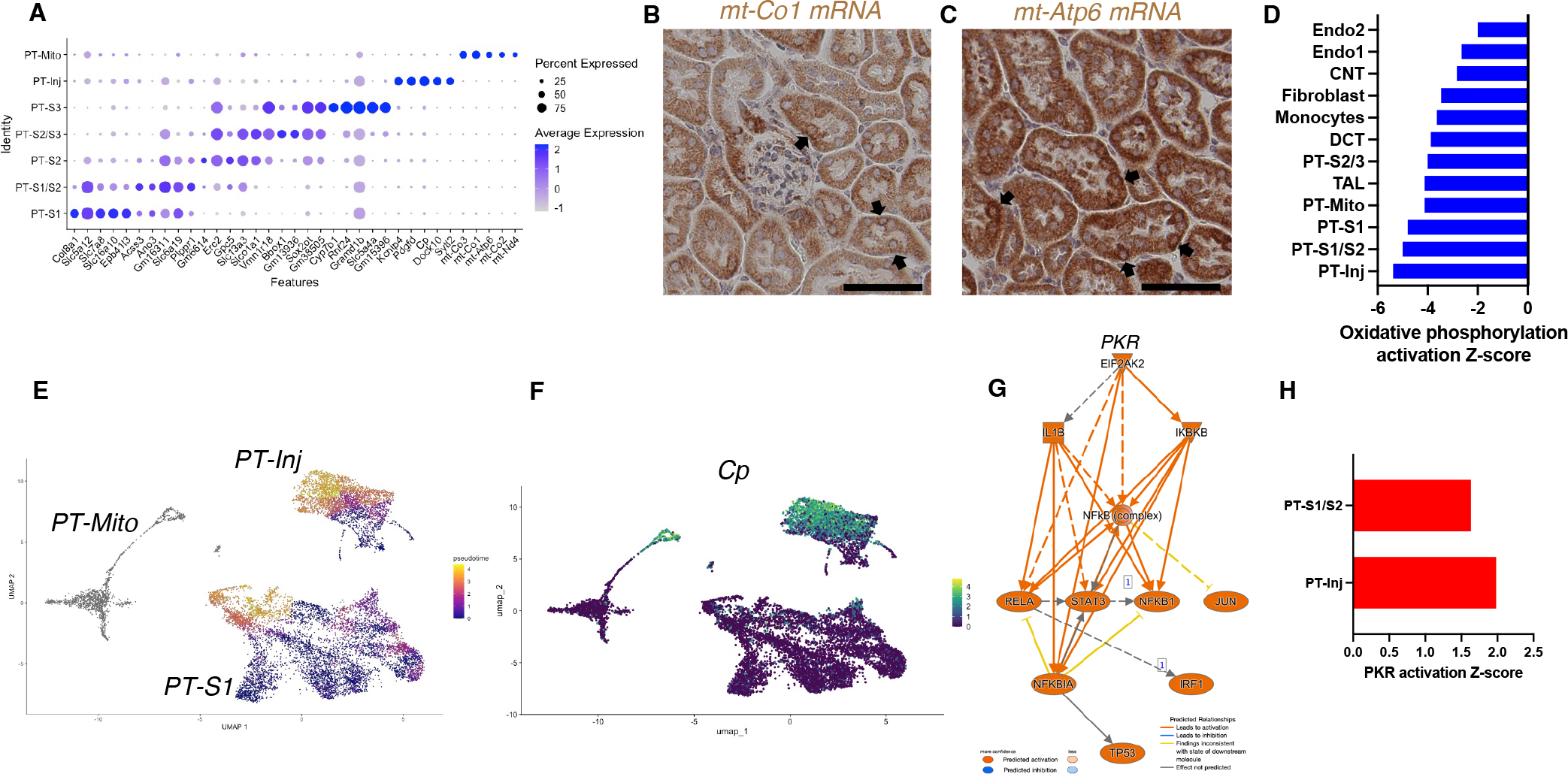
PT-Mito and PT-Inj cluster characterization. (**A**) Shown are dot plots showing the top five marker genes in each of the PT clusters (PT-S1, PT-S1/S2, PT-S2, PT-S2/S3, PT-S3, PT-Inj, PT-Mito). (**B, C**) *In situ* hybridization of *mt-Co1* and *mt-Atp6* genes showed signals inside nuclei of WT mice. (Scale bars are 50 μm) (**D**) Shown is the activation Z-score of the oxidative phosphorylation pathway by pathway analysis of each cluster comparing Tg26 vs WT mice. (**E**) Trajectory analysis results including PT-S1, PT-Mito, PT-Inj from WT and Tg26 mice. (**F**) Ceruloplasmin (*Cp*) expression in the trajectory analysis plot. (**G**) PKR downstream pathway mapping by IPA comparing Tg26 vs WT by bulk mRNA-seq data. (**H**) Activation Z-score of PKR pathway by IPA in each cluster comparing Tg26 vs WT.

As the PT-Mito cluster was observed in both WT and Tg26 kidney in similar ratios, this new cluster could be a conserved tubular cell cluster, likely not previously reported due to mitochondrial gene filtering criteria implemented in single-nucleus RNA-seq analytic pipelines. Pathway analysis of snRNA-seq data showed that oxidative phosphorylation genes were downregulated in this PT-Mito cluster in Tg26 mice, compared to wild-type mice, with a Z-score of −4.123. These genes were also downregulated in other many cell types (**Figure 4D**). These findings suggested that mitochondrial dysfunction represents a prominent mechanistic pathway that was dysregulated in majority of cell clusters in Tg26 mice, and that mitochondrial dysfunction was pronounced in the PT-Mito cluster.

### Proximal tubular cells with injury were more frequent in Tg26 mice

We identified a proximal tubular cell cluster, enriched for injury markers (PT-Inj), which was increased in cell number and percentage in Tg26 mice when compared with WT mice and C16 treated-Tg26 mice. The gene expression profile of this cluster was comparable to profiles previously-reported mouse models of acute kidney injury (AKI)^40^ and kidney fibrosis.^48^ We compared gene expression in Tg26 mice with previously reported expression of marker genes in injured proximal tubules from snRNA-seq data obtained from a mouse ischemia-reperfusion injury model (**Supplemental Figure 7A**). We found some overlap in these two models (Tg26, ischemia-reperfusion), and the higher expression of these genes in Tg26 mice was ameliorated with C16 treatment (**Supplemental Figure 7B**). This finding suggested some overlap of pathological molecular pathways between HIV-associated tubular injury and ischemic injury, especially at later stages when fibrosis appears.

Pseudotime analysis suggested that injured proximal tubular epithelial cells likely originated from PT-S1 segments, suggesting proximal tubular damage in Tg26 mouse kidneys (**Figure 4E, 4F**). Differential gene expression analysis and IPA upstream analysis of bulk RNA-seq data from Tg26 and wild type mouse kidneys showed activation of the PKR pathway (**Figure 4G**). The per cluster upstream analysis of snRNA-seq data showed the highest PKR pathway activation in PT-Inj (**Figure 4H**).

### PKR inhibition rescued Stat3 activation in Tg26 mice

PKR inhibition, acting to reduce inflammatory pathway activation, may influence other important mediators of HIVAN pathophysiology. To identify possible mediators, we used upstream analysis in IPA. We compared C16-treated WT vs WT, Tg26 vs WT, and C16-treated Tg26 vs Tg26, using differential gene expression analysis. All genes with multiple-testing adjusted p-values <0.05 were included in this analysis. An activated Z-score was calculated for each candidate upstream regulator. Candidate transcription factors were sorted in the order of triple Z-score, which was calculated by multiplying the three Z-scores (**Supplemental Figure 8**).

Stat3 and representative examples of Stat3-regulated gene expressions profiles are shown for each tissue comparison (**Figure 5A, 5B**). Interestingly, Stat3 was the most activated upstream regulator and is a well-characterized transcriptional regulator in kidney disease, including in the Tg26 mouse model.^19^ We confirmed Stat3 activation through phosphorylation by Western blot showing increased phosphorylation (**Figure 5C, 5D**) and immunohistochemistry (**Figure 5E-G**). These data suggest that PKR inhibition may be therapeutic for Tg26 kidney disease, promoting deactivation of Stat3 and downstream inflammatory pathways. Based on single-nucleus RNA-seq and IPA upstream analysis, we confirmed that Stat3 was activated in the majority of cell types in Tg26 and especially in PT-Inj (**Figure 5H**). As Stat3 translocate to mitochondria and alter cell metabolism^49^, Stat3 may also be important mitochondrial regulator in the pathogenesis of HIVAN.

**Figure. 5.**
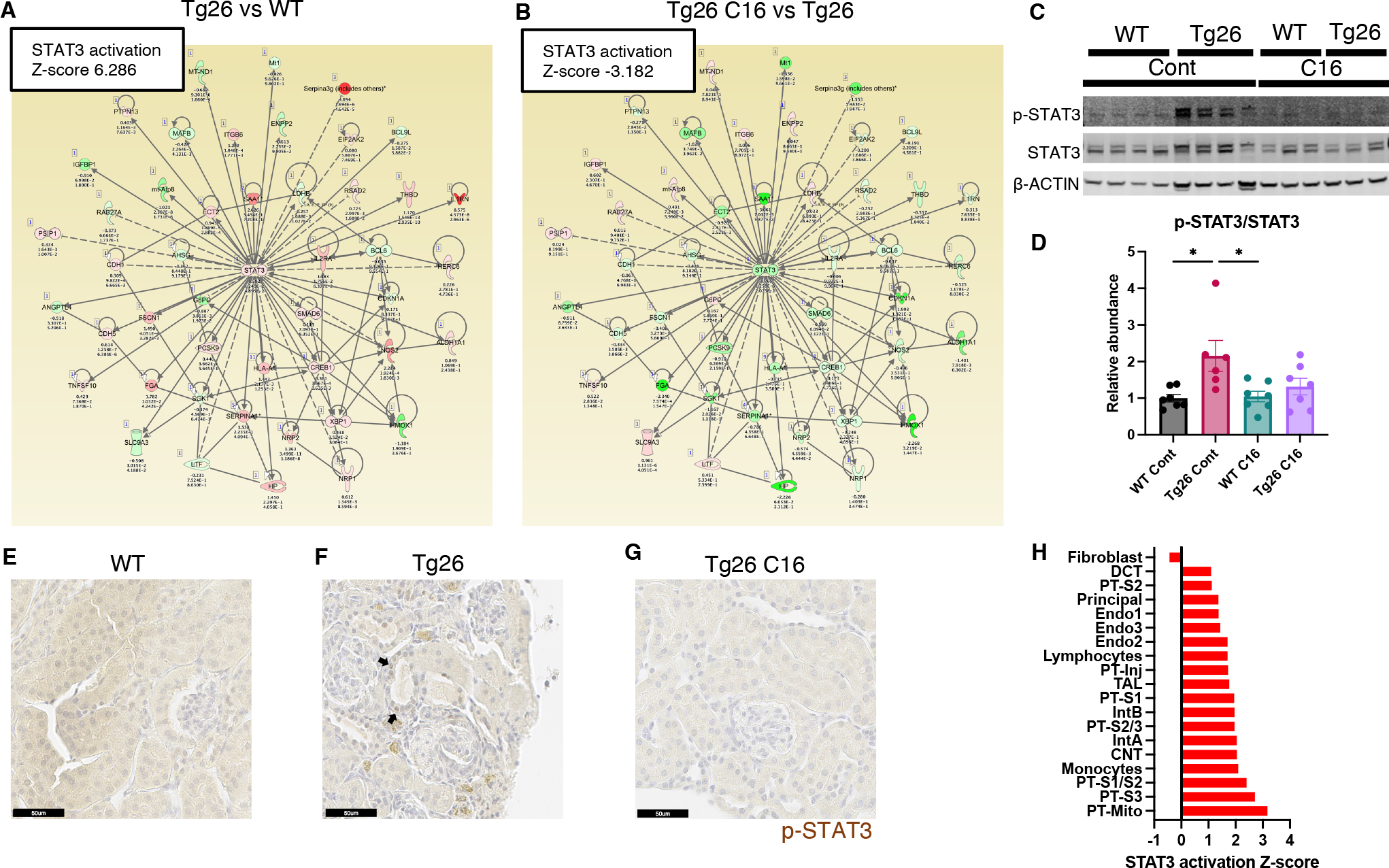
STAT3 activation downstream of PKR. (**A, B**) Shown is mapping of STAT3 regulating genes comparing Tg26 vs WT (Z-score 6.286), Tg26 C16 vs Tg26 (Z-score −3.182) by bulk mRNA-seq. Red color indicates upregulation and green color indicates downregulation, quantified as log2-fold change, p-value; and adjusted p-value by false discovery rate are shown. Solid line indicates known positive regulation, dotted line indicates known negative regulation. (**C**) Representative immunoblotting of phospho-STAT3, STAT3 and β-ACTIN. (**D**) Quantitative results of phospho-STAT3/STAT3 by immunoblotting. (one-way ANOVA; *, P<0.05) (**E-G**) Phospho-STAT3 immunostaining of mouse kidneys is shown (Scale bars are 50 μm). Arrows indicate p-Stat3 detection in injured tubular cells. (**H**) STAT3 activation Z-score by upstream regulator analysis comparing Tg26 vs WT in each cluster by snRNA-seq.

### PKR inhibition restored reduced mitochondrial respiration capacity in Tg26

These data shown indicated that mitochondrial-expressed gene transcription was inhibited in the kidney cortex of Tg26 mice and was rescued by PKR inhibition. To investigate mitochondrial functions in kidney, we prepared enriched glomerular and proximal tubular tissue extracts. These were tested for mitochondrial respiratory capacity using the Seahorse extracellular flux analyzer. The results showed reduced maximum respiration and spare respiratory capacity in Tg26 compared to wild-type mice (**Figure 6A-C**). Reduced respiratory capacity in Tg26 mice was rescued by C16 treatment, suggesting that PKR inhibition restored impaired mitochondrial respiration in Tg26 mice. Similarly, proximal tubular tissue *ex vivo* also showed lower maximum respiration and less spare respiratory capacity in Tg26 mice compared to wild-type mice (**Figure 6D-F**); both parameters were normalized by C16 treatment. Thus, PKR inhibition rescues mitochondrial dysfunction in both glomerular and proximal tubular cells.

**Figure. 6.**
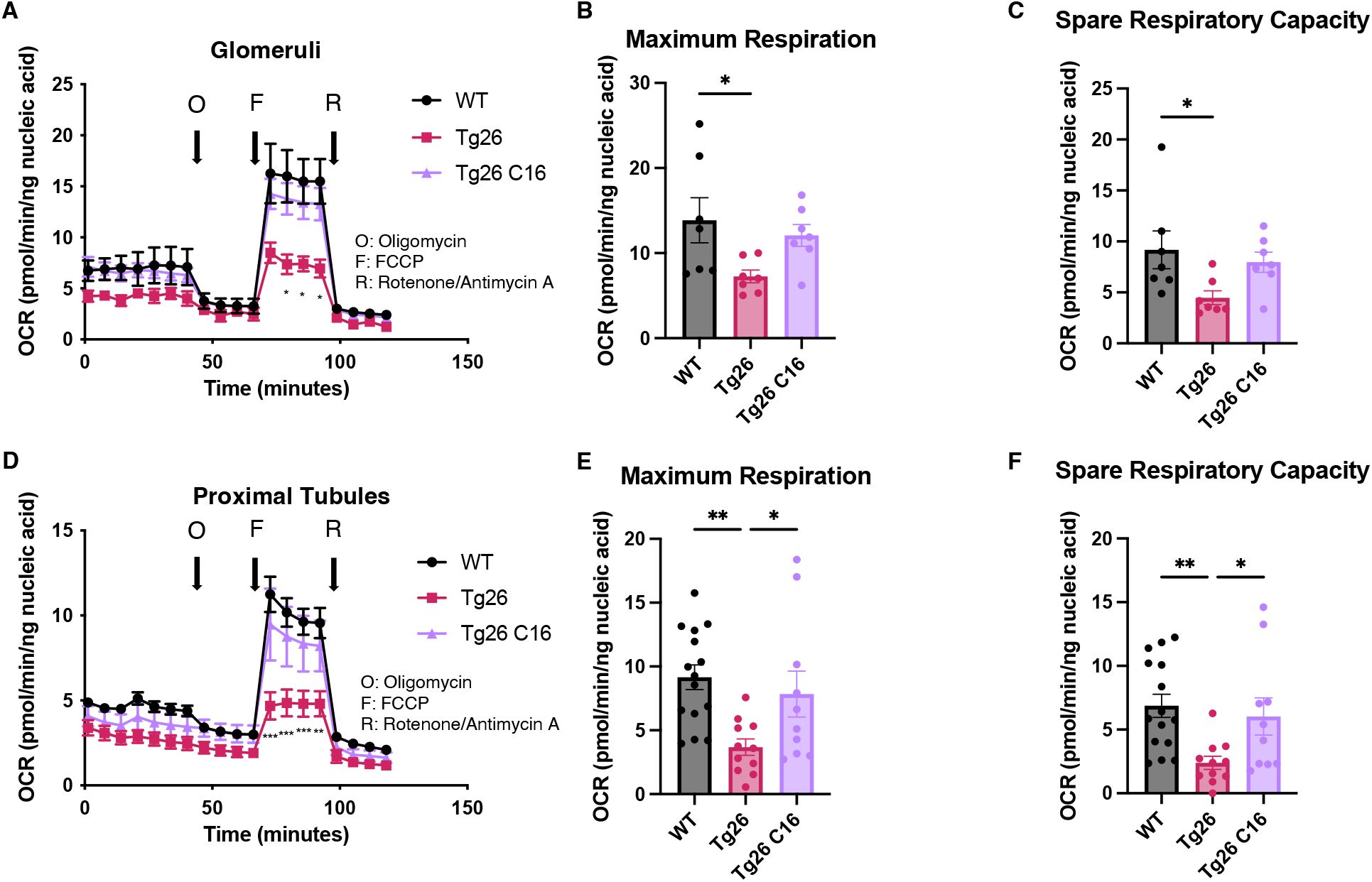
PKR inhibition with C16 reverses mitochondrial dysfunction in Tg26 glomeruli and proximal tubules. (**A**) Shown are oxygen consumption rate (OCR) measurements during cell mitochondrial stress testing, using extracted glomeruli from WT, Tg26, and C16 Tg26 kidney tissue. (**B**) Maximum respiration rate was calculated by OCR measurements of glomeruli. (**C**) Spare respiratory capacity was calculated by OCR measurements of glomeruli. (**D**) Shown are oxygen consumption rate (OCR) measurements during cell mitochondrial stress testing, using extracted proximal tubules from WT, Tg26, and C16 Tg26 kidney tissue. (**E**) Maximum respiration rate was calculated by OCR measurements of isolated proximal tubules. (**F**) Spare respiratory capacity was calculated by OCR measurements of isolated proximal tubules. (One-way ANOVA; *, P<0.05; **, P<0.01)

### Podocytes in Tg26 mice showed high HIV-1 gene expression with podocyte loss

Although the Tg26 mouse is a well-characterized model of HIVAN, expression levels of each of the HIV-1 genes in single kidney cells has not been previously reported. Using single-nucleus RNA-seq, we annotated expression of each HIV-1 gene, with the aim to quantify HIV-1 gene expression at single cell resolution (**Figure 7A**). Among all kidney cell types, HIV-1 genes were expressed at high levels in podocytes, consistent with the notion that HIV-1 gene expression is particularly injurious to podocytes. HIV-1 gene expression data from this study are consistent with the current understanding that *vpr* and *nef* are main contributors to the pathogenesis of HIVAN. Absolute podocyte loss in Tg26 mice, and podocyte recovery with C16 treatment, were also confirmed by p57 staining and podocyte estimation using the podometric analysis implemented in PodoCount (**Figure 7B-E**).

**Figure. 7.**
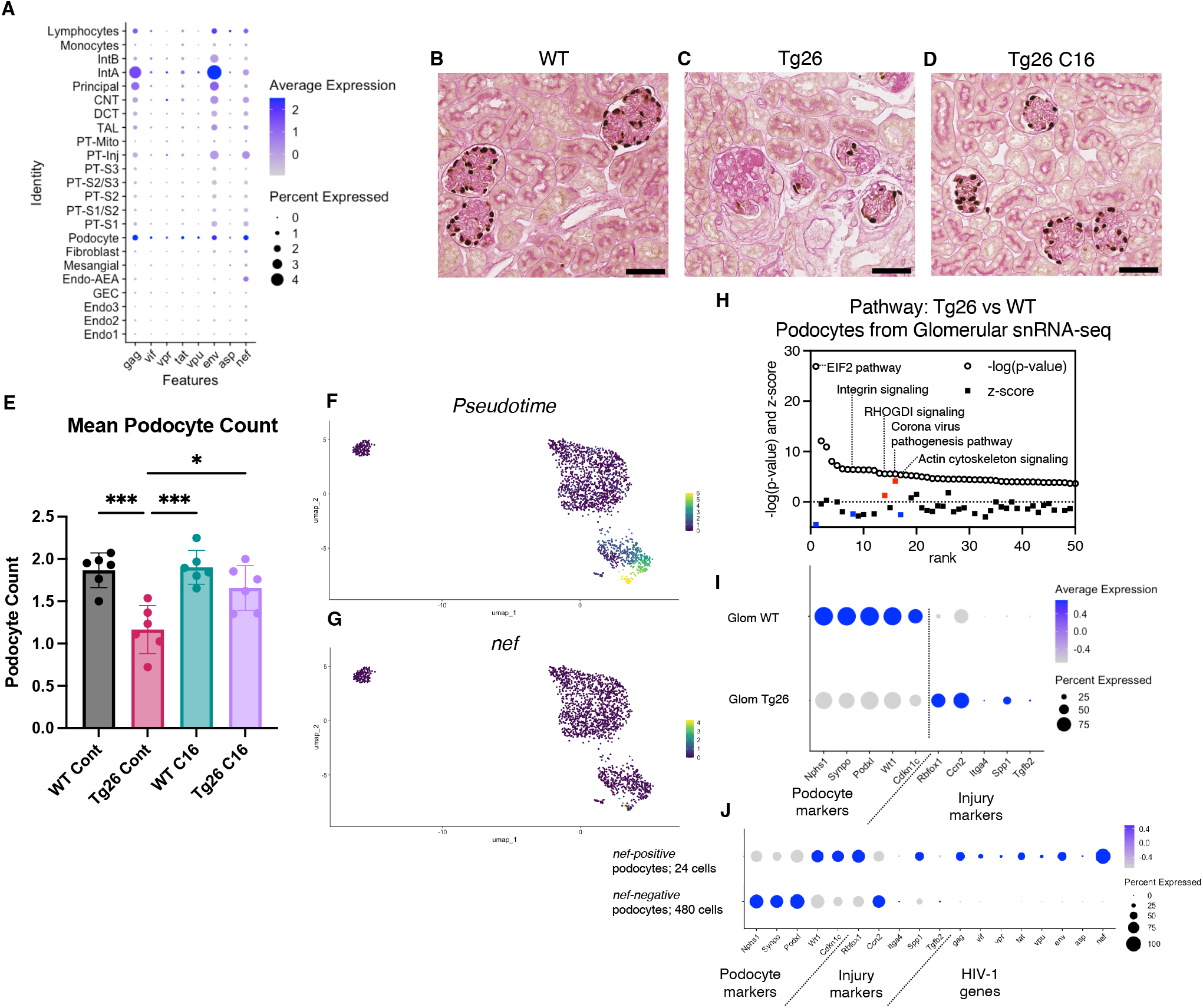
HIV-1 gene expression causes Tg26 podocytes dedifferentiation. (**A**) Shown is dot plot demonstrating HIV-1 gene expression levels in each cluster detected by snRNA-seq. (**B-D**) p57 staining of kidney showing podocyte loss and dedifferentiation. (Scale bars are 50 μm) (**E**) Podocount analysis showed podocyte loss in Tg26 and was rescued by C16. (One-way ANOVA; *, P<0.05; ***, P<0.001) (**F**) Trajectory analysis of podocytes by snRNA-seq data from WT, Tg26 glom samples. (**G**) Trajectory map showing *nef* expression. (**H**) Pathway analysis results by IPA comparing Tg26 vs WT using glomerular snRNA-seq data from podocyte cluster. (**I**) Dot plot showing podocyte marker genes and representative differentially expressed genes in podocytes by glomerular snRNA-seq. (**J**) Dot plot comparing expression of representative genes in glomerular Tg26 podocytes between *nef*-positive and *nef*-negative podocytes.

### Podocytes in Tg26 mice showed dedifferentiation and dysregulation of cytoskeleton-related pathways

Since podocytes in Tg26 mice expressed the HIV-1 genes *vpr* and *nef*, we investigated related pathways that were dysregulated in podocytes. The yield of podocytes from kidney cortex samples was relatively low and, therefore, we used snRNA-seq data from isolated glomeruli from wild-type and Tg26 mice, which enriched podocytes. Pseudotime analysis of these data showed progression in transcripts from wild-type podocytes to the majority of Tg26 podocytes, showing *nef* expression in most injured Tg26 podocytes (**Figure 7F, 7G**). Differential expression analysis and pathway analysis suggested that in Tg26 mice, Rho protein dissociation inhibitors (RHOGDI) signaling and corona virus pathogenesis pathways were activated, while integrin and actin cytoskeleton signaling pathways were deactivated (**Figure 7H**).

These findings are consistent with previous reports and with a common conceptual model in which podocyte dedifferentiation starts with cytoskeletal changes and progresses to cell detachment from the glomerular tuft.^50–52^ We confirmed dedifferentiation of podocytes in Tg26 mice by showing down-regulation of podocyte markers and reversal or prevention of dedifferentiation in Tg26 mice treated with C16 (**Figure 7I**). This was also demonstrated histologically by p57 staining, which was lost in Tg26 and then regained following C16 treatment (**Figure 7B-D**). We also found increased expression of *Ccn2* (encoding cellular communication network factor 2), an epithelial–mesenchymal transition (EMT) marker, in Tg26 podocytes. This is consistent with the signatures that we previously reported in urine podocytes in a single-cell clinical study of FSGS patients.^53^ We also observed upregulation of *Rbfox1*, encoding RNA-binding fox-1 homolog-1, as candidate disease marker in Tg26 podocytes (**Figure 7I**). Rbfox1 also regulates key neuronal functions.^54,55^

Expression of integrin subunits *Itga4* and of *Spp1* (encoding osteopontin) and *Tgfb2* (encoding transforming growth factor β2) was also upregulated in Tg26 podocytes, indicating activation of cell adhesion and pro-fibrotic processes (**Figure 7I**). We compared gene expression in *nef*-positive podocytes (24 cells) and *nef-negative* podocytes (480 cells) from Tg26 glomerular data, to confirm the association between HIV-1 gene expression and transcriptomic changes. We found lower expression of canonical podocyte marker genes, including *Nphs1*, *Synpo*, together with higher expression of *Rbfox1* and *Spp1* described above (**Figure 7J**).

### Tg26 kidney cells showed active cell-cell interaction

To investigate pathologic cell-cell interactions in Tg26 kidneys and the ameliorative effect of PKR inhibition, we performed cell-cell interaction analysis using the snRNA-seq data. The expression heatmap suggested changes in cell-cell interaction in each sample at the resolution of each identified cell cluster (**Supplemental Figure 9A-C**). We sought candidate cell types that might contribute to Tg26 pathology and found candidate ligand-receptor pairs (**Figure 8A-C**). For example, platelet derived growth factor D (PDGF-D) was upregulated in PT-Inj in Tg26 mice and was downregulated by C16 treatment **(Figure 8D**). Further, PDGF-D may interact with PDGFR-B in fibroblasts. Immunohistochemistry demonstrated the presence of PDGF-D in the vicinity of dilated tubules (**Figure 8E, 8F**). As PDGF-D is known to be induced by STAT3, these findings identified a potential fibrogenic pathway triggered by PKR activation in PT-Inj.

**Figure. 8.**
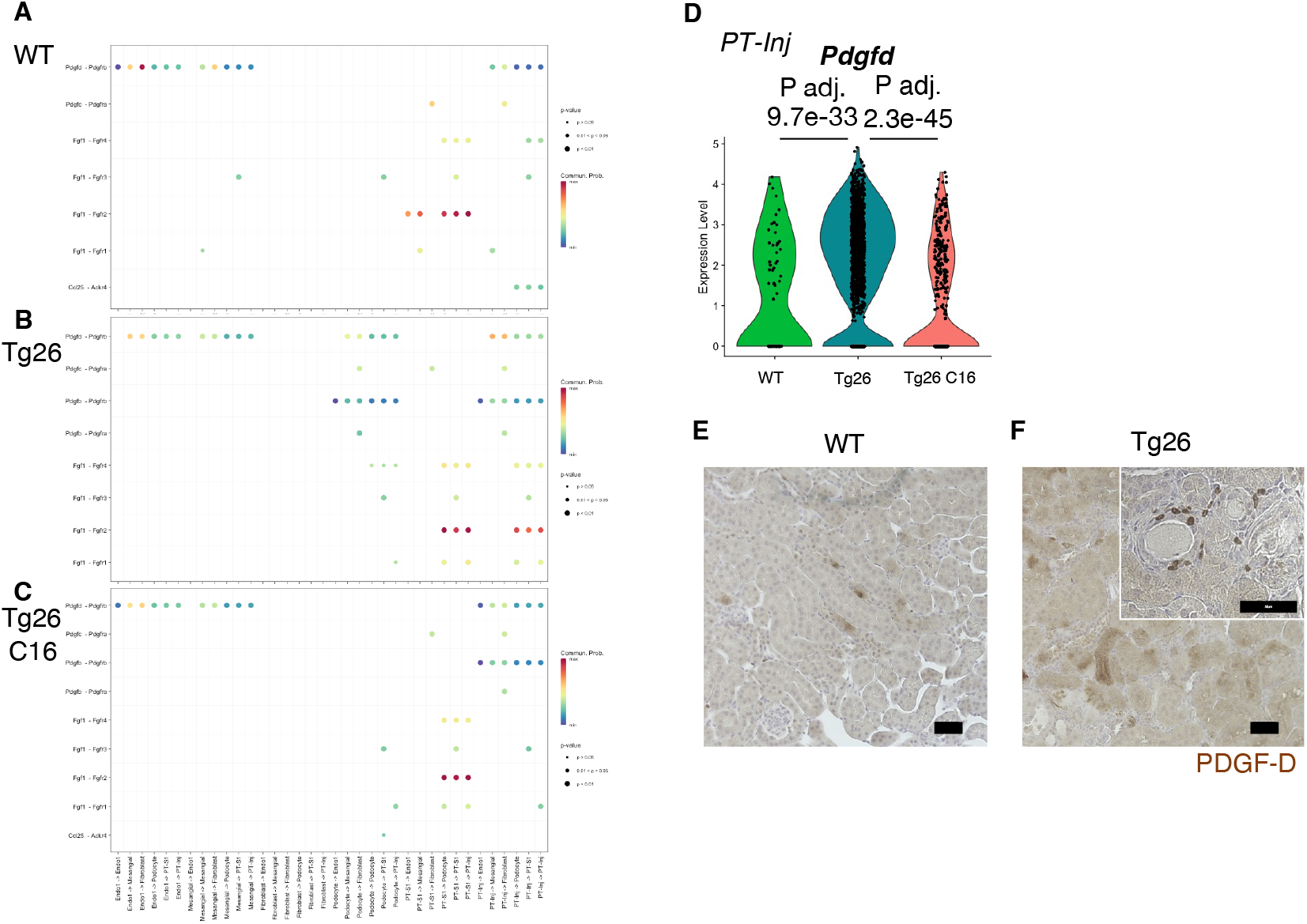
Cell-cell interaction analysis shows activated ligand-receptor interaction: PDGF-D-PDGFR-B pathway in Tg26. (**A-C**) Shown are dot plots depicting results from cell-cell interaction analysis of WT, Tg26, C16 Tg26 snRNA-seq data. (**D**) Violin plots showing *Pdgfd* expression levels in the PT-Inj cluster. (**E, F**) PDGF-D immunostaining of mouse kidneys is shown. (Scale bars are 50 μm)

## Discussion

This work, applying combined analysis of single-nucleus RNA-seq and bulk RNA-seq data, highlight the role of PKR activation in the Tg26 HIVAN mouse model and show mitochondrial dysfunction to be one of the most dysregulated pathways. Mitochondrial dysfunction in Tg26 mouse glomeruli and myocytes has been previously reported.^24,56^ The present study demonstrated by single-nucleus RNA-seq and Seahorse bioenergetics assay that mitochondrial dysfunction is induced in various cell types in Tg26 kidney, especially proximal tubular cells and endothelial cells. It is likely that HIVAN tubular pathology is mediated, at least in part, by mitochondrial dysfunction in proximal tubular cells, and that glomerular mitochondrial dysfunction is mainly confined to endothelial cells. Further, PKR activation may be one of several processes that contributes to mitochondrial dysfunction in HIVAN. A recent report using tubular cell-specific Vpr overexpression mouse model has shown that mitochondrial dysfunction is implicated in tubular injury. Our findings are congruent with this report.^57^

Further, single-nucleus RNA-sequencing of Tg26 kidney cortex identified a novel mitochondrial gene-enriched proximal tubular cell population (PT-Mito). This cell population had not been previously recognized, likely due to filtering criteria routinely employed to reduced mitochondrial genome-encoded transcript levels in single-cell/single nucleus datasets. It has been well established that a high percentage of mitochondrial transcripts captured in single-cell RNA-seq studies suggests stressed cells, and these cells are typically excluded from analysis. However, in single-nucleus RNA-seq data sets, mitochondrial gene percentage is generally low following the nuclear purification. Most nuclei captured have mitochondrial transcripts levels of <1% of the total transcripts captured. Therefore, we did not filter out mitochondrial transcripts entirely from the analysis but excluded nuclei with >20% mitochondrial transcripts^58^ By not filtering out mitochondrial genes in the analysis, we found dysregulation of these transcripts, providing clues to pathogenesis.

We further demonstrated by ISH the presence of mitochondrial transcripts in nuclei whose cells were identified in single-nuclei analysis of the same tissue samples. Although we cannot completely exclude the possibility of having captured some mitochondria during nuclear preparation, we confirmed expression of nuclear genes, such as *Gpx1* and *Gpx3*, which were also highly expressed in this PT-Mito cluster. A relatively high abundance of mitochondrial transcripts in the PT-Mito cluster may indicate the existence of mitochondrial transcripts that were transported into nuclei. This PT-Mito cluster might have remained previously undetected because of the similarity of its transcripts with that of those other proximal tubules and the lack of cell type-specific markers. Considering the high mitochondrial gene expression levels and corresponding mitochondrial pathway dysregulation in Tg26 mice, this PT-Mito cluster may represent the most metabolically active cells, which are also the most highly vulnerable cell type in proximal tubules. Nevertheless, considering the downregulation of both nuclear and mitochondrial encoded genes involved in oxidative phosphorylation; mitochondrial dysfunction likely plays a major role in the pathology of HIVAN.

Another novel finding arose from investigating HIV-1 gene expression in the Tg26 mouse. Transcripts for all transgene-encoded HIV-1 genes were detected in all cell types examined, albeit some transcripts were present at low expression levels. This included transcripts for *nef* and *vpr* that are, according to current understanding, the main contributors to HIVAN.^59,60^ Podocytes showed the highest levels of transgene expression, which is consistent with the prominent pathology observed in podocytes. This finding suggests a shared but unknown mechanism in virus-related nephropathies with podocyte damage. One possible explanation for podocytes not showing overt mitochondrial gene dysregulation despite high HIV-1 gene expression (compared to proximal tubular cells and endothelial cells) is a possible tighter regulation of mitochondrial gene expression in podocytes, as suggested by Li et al.^61^ With regard to S phase-specific genes^62^, we did not find a consistent expression change in cells including podocytes from the glomerular sample (**Supplemental Figure 3G**).

Further, we identified a putative activated pathway involving PKR - STAT3 - PDGF-D - PDGFR-B in injured proximal tubules. The PKR-STAT3-PDGF signaling cascade has been reported in PKR-null mouse embryonic fibroblasts.^63^ PDGF-D and PDGFR-B contribute to fibrosis in glomeruli and the tubulointerstitium in experimental animal studies ^64,65^. Inhibiting this pathway may offer an avenue to reduce kidney injury in HIVAN. Further studies will be needed to confirm whether this mechanism is shared with human HIVAN or other RNA virus-associated kidney diseases.

The present study has limitations. First, the Tg26 mouse model involves a partial HIV-1 transgene that may not recapitulate all aspects of clinical HIVAN. Second, gene expression changes after C16 treatment may include changes secondary to the attenuated renal injury, in addition to the direct effect of C16. Third, we acknowledge possibility of a non-specific effect of C16 as an inhibitor of PKR.^66–68^

In conclusion, by combining single-nucleus RNA-seq and bulk RNA-seq analysis, we identified mitochondrial dysfunction as the central mechanism for proximal tubule injury in the Tg26 HIVAN mouse model. This process that was largely reversed by treatment with the PKR inhibitor C16. Further studies of HIVAN-associated mitochondrial dysfunction may lead to targeted therapeutics.

## Author Contributions

TY, KO, JBK conceived the study design. TY conducted mouse experiments with support by SS. TY analyzed bulk RNA-seq data. TY and YY conducted single-nuclear RNA-seq capture. TY and KZL analyzed single-nuclear RNA-seq data. YZ and CAW supported sequencing at FNLCR/NCI. AZR assessed pathological quantification. BS, TY, VT and PS conducted podocyte morphometry. KM, BJ, XW conducted OXPHOS complex Western blot. TY, JBK drafted the manuscript and all the authors contributed for edits. The order of the co-first authors was based on when they started on the project and the relative contribution to the authoring of the original version of the manuscript.

## Acknowledgement

We thank the Sequencing Facility and Bioinformatics Group (Frederick National Laboratory for Cancer Research (FNLCR), NCI, NIH) for sequencing and informatics support, Drs. Joon-Yong Chung and Stephen M. Hewitt (NCI/NIH) for whole slide scanning, Maria Campos (NEI/NIH) for pathological service, Dr. Daria Ilatovskaya (Medical University of South Carolina) for suggestion of Seahorse assay, and Drs. Mark A. Knepper (NHLBI, NIH), Gregory G. Germino (NIDDK, NIH) and Michael J. Ross (Albert Einstein College of Medicine) for scientific suggestions and supports, Dr. Jurgen Heymann for critical manuscript review. This work utilized the computational resources of the NIH HPC Biowulf cluster. (http://hpc.nih.gov) Part of this work was presented at American Society of Nephrology Kidney Week 2020, 2021. The content of this publication does not necessarily reflect the views or policies of the Department of Health and Human Services, nor does mention of trade names, commercial products, or organizations imply endorsement by the U.S. Government.

## Data Availability

Original data files and count tables have been deposited in GEO (GSE205060). Other data are available from the authors upon request.

