## Supplemental Materials for "PKR activation-induced mitochondrial dysfunction in HIV-transgenic mice with nephropathy"

Running Title: PKR activation in murine HIV-associated nephropathy

Teruhiko Yoshida[1*#], Khun Zaw Latt[1#], Avi Z. Rosenberg[2], Briana A. Santo[3], Komuraiah Myakala[4], Yu Ishimoto[5], Yongmei Zhao[6], Shashi Shrivastav[1], Bryce A. Jones[4], Xiaoping Yang[2], Xiaoxin X. Wang[4], Vincent M. Tutino[3], Pinaki Sarder[3,7], Moshe Levi[4], Koji Okamoto[1,8], Cheryl A. Winkler[6], Jeffrey B. Kopp[1]

[1] Kidney Disease Section, Kidney Diseases Branch, NIDDK, NIH, Bethesda, MD

[2]Department of Pathology, Johns Hopkins Medical Institutions, Baltimore, MD

[3]Department of Pathology and Anatomical Sciences, Jacobs School of Medicine & Biomedical Sciences, University at Buffalo, Buffalo, NY

[4]Department of Biochemistry and Molecular & Cellular Biology, Georgetown University, Washington, DC

[5]Polycystic Kidney Disease Section, Kidney Diseases Branch, NIDDK, NIH, Bethesda, MD

[6]Frederick National Laboratory for Cancer Research, NCI, NIH, Frederick, MD

[7]College of Medicine, University of Florida, Gainesville, FL

[8]Nephrology Endocrinology and Vascular Medicine, Tohoku University Hospital, Sendai, Japan

[#] These authors contributed equally to this manuscript.

*Corresponding author:* Teruhiko Yoshida, MD PhD;

*address:* 10 Center Dr. 3N104, Bethesda, MD 20892-1268

### Table of Contents

**Supplemental Methods**

**Supplemental Figures**

Supplemental Figure 1. Immunofluorescent images showing pPKR

Supplemental Figure 2. Pre-processing of single-nuclear RNA-seq data

Supplemental Figure 3. Additional plots of single-nuclear RNA-seq data

Supplemental Figure 4. Additional data about mitochondria and S phase score of podocytes

Supplemental Figure 5. Additional *in situ* hybridization images

Supplemental Figure 6. PT-Mito cluster detection of publicly available human kidney single-nuclear RNA-seq data (GSE131882)

Supplemental Figure 7. Single-nuclear RNA-seq data comparison with ischemic reperfusion injury model

Supplemental Figure 8. Activation Z-scores from bulk RNA-seq data

Supplemental Figure 9. Additional cell-cell interaction analysis plot

### Supplemental Methods

*Mouse chemistry measurements*

Plasma creatinine was measured by isotope dilution LC-MS/MS at The University of Alabama at Birmingham O'Brien Center Core C (Birmingham, AL). Urine albumin levels were measured using Albuwell M ELISA kits (Ethos Biosciences, Newtown Square, PA). Urine creatinine concentrations were measured using the Creatinine Companion kit (Ethos Biosciences, Newtown Square, PA). Urinary NGAL and KIM-1 were measured by the Mouse Lipocalin-2/NGAL DuoSet ELISA and Mouse TIM-1/KIM-1/HAVCR Quantikine ELISA Kit (R&D Systems, Minneapolis, MN).

*Glomerular and proximal tubule enrichment method*

Mice were anesthetized using 2, 2, 2-trimbromoethanol (Avertin) and the abdominal aorta and vena cava were exposed. After clipping the abdominal aorta distal to renal artery bifurcation, a catheter was inserted into the aorta proximal to an incision above iliac artery bifurcation. After clipping the celiac trunk, superior mesenteric artery, and thoracic aorta proximal to renal arteries bifurcation, a small incision was made in the renal vein to perfuse both kidneys with 1 ml of PBS. Next, kidneys were perfused twice with 10 ul of Dynabeads M-450 Tosylactivated (#14013, Thermo Fisher Scientific, Waltham, MA), to facilitate isolation of glomeruli.

Kidneys were immediately collected, decapsulated, and placed into Hanks Balanced Salt Solution (HBSS) medium on ice. Kidneys were minced using razor blades and enzymatically digested in 1 mL of HBSS containing 4 mg collagenase A (#10103586001, Sigma, Darmstadt, Germany) and 40 µl DNase I recombinant (E04716728001, Sigma, Darmstadt, Germany) for 30 mins at 37 °C with shaking at 1500 rpm. Glomerular samples were collected after filtration through a 100 μm strainer, followed by magnetic separation (MPC-S, DYNAL) and three PBS washes. Proximal tubular samples were collected from the non-glomerular supernatant of the first magnetic separation step. Proximal tubules were isolated by centrifugation through 31% Percoll (17089102, Cytiva, Marlborough, MA)-PBS centrifugation, followed by an additional wash and centrifugation with PBS, following a published protocol.[1,2]

*Seahorse Extracellular Flux Assay*

Seahorse 96-well assay plates (Agilent, Santa Clara, CA) were pre-coated twice with 20 μl/well of 0.01% poly-L-lysine solution (P4707, Sigma, Darmstadt, Germany) and washed twice with PBS, 200 μl/well. Glomerular or proximal tubular samples were plated with EGM-2 medium (CC-3162, Lonza, Walkersville, MD) and placed in a $CO_2$-incubator for 30 minutes for the attachment.

Seahorse XF RPMI medium, pH 7.4 (103576-100, Agilent, Santa Clara, CA), was used for glomerular samples and Seahorse XF DMEM medium, pH 7.4 (103575-100, Agilent, Santa Clara, CA), was used for proximal tubular samples. Media were supplemented to reach a final concentration of 10 mM glucose, 1 mM sodium pyruvate, and 2 mM L-glutamine. Reagent concentrations used were 2 μM oligomycin, 2 μM FCCP, 0.5 μM rotenone and 0.5 μM antimycin A (103015-100, Agilent, Santa Clara, CA).

Seahorse Mito Stress Tests were conducted as described.[3] Cells were incubated in a $CO_2$-free incubator for 30 minutes, after replacement of medium. Data were normalized by total nucleic acid content measured by CyQUANT Cell Proliferation (C7026, Thermo Fisher Scientific, Waltham, MA) and analyzed by Wave 2.6.1 (Agilent, Santa Clara, CA).

*Mitochondrial copy number measurements*

Determination of mitochondrial copy number of mouse kidney tissues was conducted following a published method.[4] Genes encoding 16S rRNA and *Nd1*were measured as mitochondrial DNA (mtDNA) and the *Hk2* gene was measured as nuclear DNA (nDNA).The mtDNA/nDNA ratios in mouse tissues were quantified by SYBR green assay using QuantStudio 6 (Thermo Fisher Scientific, Waltham, MA).

*In situ hybridization (ISH) of mouse kidneys*

Chromogenic *in situ* detection of RNA was performed using RNAscope (Advanced Cell Diagnostics, Biotechne, Minneapolis, MN). Briefly, 5 µm tissue sections were de-paraffinized, boiled with RNAscope target retrieval reagent for 15 min and protease digested at 40 °C for 30 min, followed by hybridization for 2 h at 40 °C with RNA probe Mm-mt-Co1, Mm-mt-Atp6 (catalog # 517121, 544401). RNA probe-Mm-PPIB (catalog # 313911) and RNA Probe-DapB (catalog # 310043) were used for positive and negative control, respectively. Specific probe binding sites were visualized using RNAscope 2.5 HD Reagent Kit (catalog # 322310).

*Immunohistochemistry in mouse kidneys*

Mouse kidney tissues were fixed with 10% buffered formalin for 24 hours, embedded in paraffin, and sectioned at 4-5 µm. The sections were deparaffinized/rehydrated and antigen retrieval was performed by heating in citrate-buffered medium for 15 min in a hot water bath. Tissues were blocked with 2.5% normal horse serum for 20 mins. Sections were incubated for 1 h at room temperature with primary antibody against phospho-Stat3 (Tyr705) (#9145, 1:100 dilution, Cell Signaling, Danvers, MA), and platelet-derived growth factor (PDGF)-D (ab181845, Abcam, 1:100 dilution, Cambridge, UK). Sections were processed following ImmPRESS HRP Universal Antibody (horse anti-mouse/rabbit IgG) Polymer Detection Kit and ImmPACT DAB EqV

Peroxidase (HRP) Substrate (Vector Laboratories, Burlingame, CA) protocol, and counter stained with hematoxylin.

*Estimation of glomerular podocyte count*

PodoCount[5], a computational tool for whole slide podocyte estimation from digitized histologic sections, was used to detect, enumerate, and characterize podocyte nuclear profiles in the glomeruli of immunohistochemically labeled (IHC-labeled) murine kidney sections. Formalin-fixed, paraffin embedded tissues (2 μm thickness) were IHC-labeled for $p57^{kip2}$, a marker of podocyte terminal differentiation (ab75974, Abcam, Cambridge, UK), and detected with horse radish peroxidase (RU-HRP1000, Diagnostic BioSystems, Pleasanton, CA) and diaminobenzidine chromogen substrate (BSB0018A, Bio SB, Santa Barbara, CA). A periodic acid-Schiff post-stain was applied without hematoxylin counterstain. The tool uses a combination of stain deconvolution, digital image processing, and feature engineering to compute histologic podometrics[6] with correction for section thickness[7]. In this study, PodoCount was used to assess mean glomerular podocyte count per mouse.

*Single-nucleus RNA-seq Analysis: Cell cycle analysis*

Cell cycle analysis was performed by converting cell cycle marker genes from Tirosh et al.[8] to mouse orthologs.

*Single-nucleus RNA-seq Analysis: Comparative analysis with mouse ischemic-reperfusion injury model data reported by Kirita et al.*

A List of marker genes in injured proximal tubules (NewPT1 and NewPT2) reported by investigators using single-nucleus RNA-seq[9] in a mouse ischemic-reperfusion injury model were obtained from the Kidney Interactive Transcriptomics website (https://humphreyslab.com/SingleCell/) for comparison with our data.

*Single-nucleus RNA-seq Analysis: PT-Mito cluster in human diabetic kidney disease study reported by Wilson et al.*

The data from the single-nucleus RNA sequencing study of human diabetic kidney disease (diabetic kidney disease n=3; control n=3) reported by Wilson et al.[10] was downloaded from the Gene Expression Omnibus (GEO) using accession number GSE131882. The data was analyzed using Seurat package version 5.0.1 without removing the mitochondrial gene transcripts.

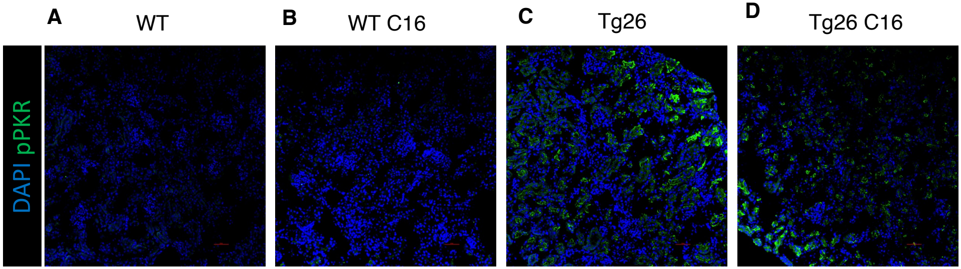


**Supplemental Figure. 1 Immunofluorescent images showing pPKR**
(**A-D**) Immunofluorescent images showed PKR activation by detecting pPKR in Tg26 mouse kidney. pPKR was inhibited by C16 treatments.

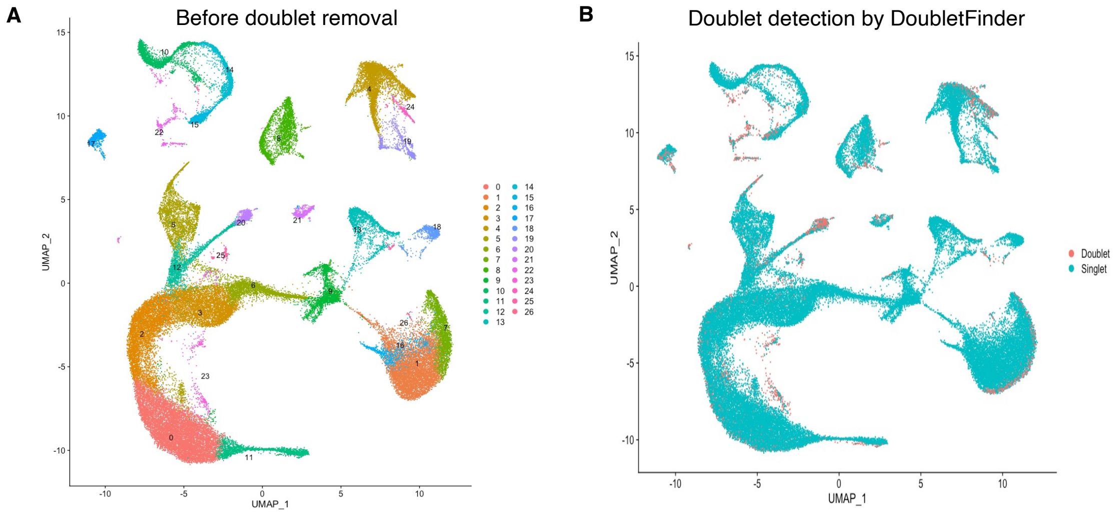


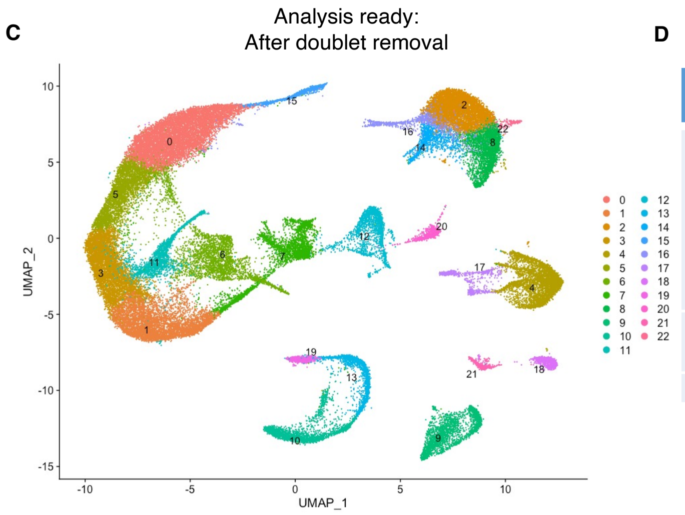


D

| Origin | Experimental Group | Sample Name | Nuclei numbers |
|---|---|---|---|
| Kidney Outer Cortex | WT | WTa | 7992 |
| | WT | WTb | 7510 |
| | Tg26 | Tg26a | 6191 |
| | Tg26 | Tg26b | 6508 |
| | Tg26 C16 | C16_Tg26a | 8821 |
| | Tg26 C16 | C16_Tg26b | 7482 |
| Glomeruli-enriched | WT | G_WT | 7224 |
| | Tg26 | G_Tg26 | 5248 |
| Total | | | 56976 |

**Supplemental Figure. 2 Pre-processing of single-nuclear RNA-seq data**
(**A**) UMAP plot of single-nuclear RNA-seq data before doublet removal from 8 samples, 57,061 cells , showing 27 clusters.
(**B**) UMAP plot showing detected doublet by DoubletFinder
(**C**) UMAP plot after doublet removal, showing 23 clusters, 56,976 nuclei for the analysis.
(**D**) Breakdown of nuclei numbers from each sample showed comparable numbers of nuclei analyzed.

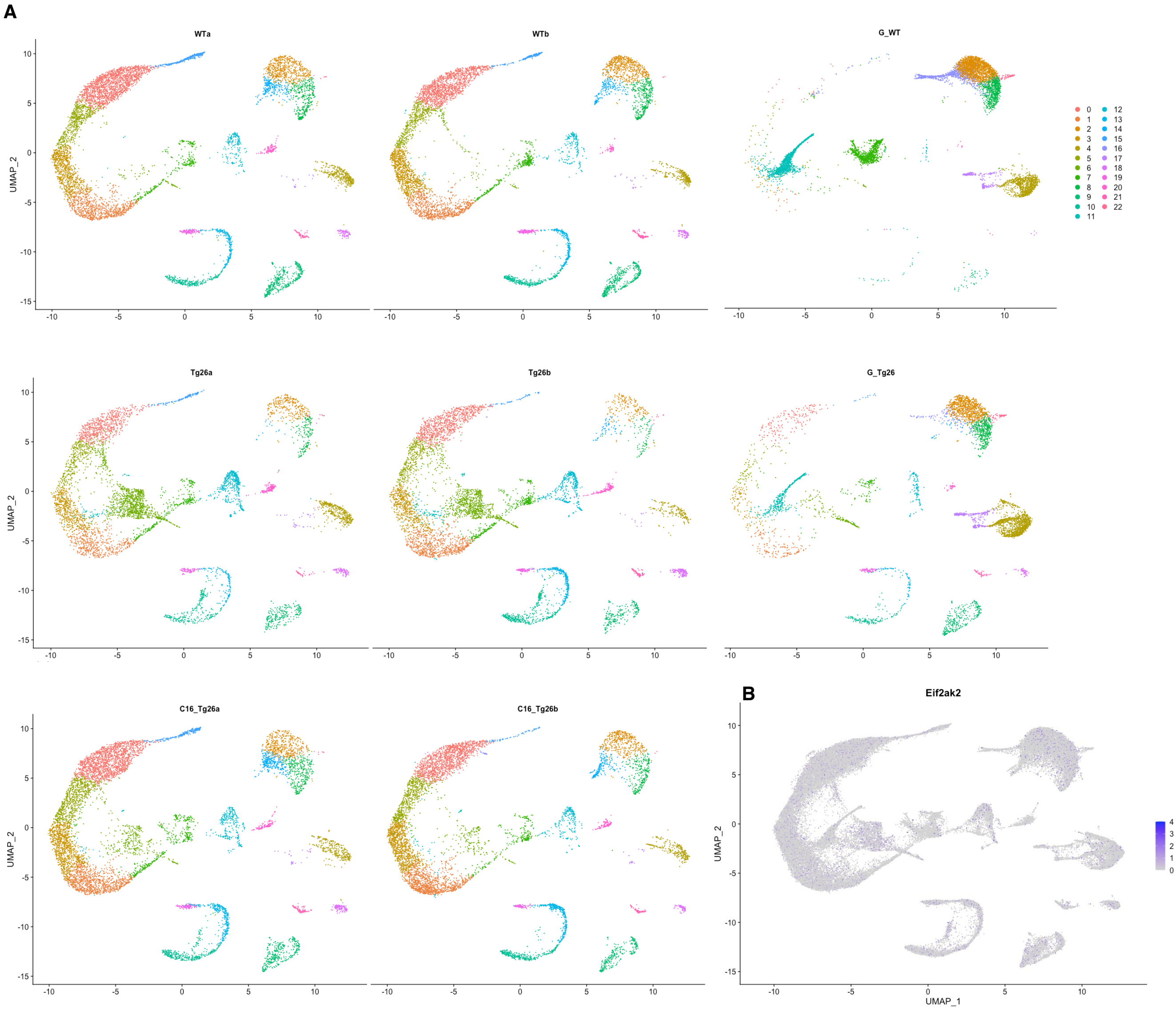


**Supplemental Figure. 3 Additional plots of single-nuclear RNA-seq data**
(**A**) UMAP plot of single-nuclear RNA-seq data from each 8 samples.
(**B**) Feature plot showing *Eif2ak2* expression.

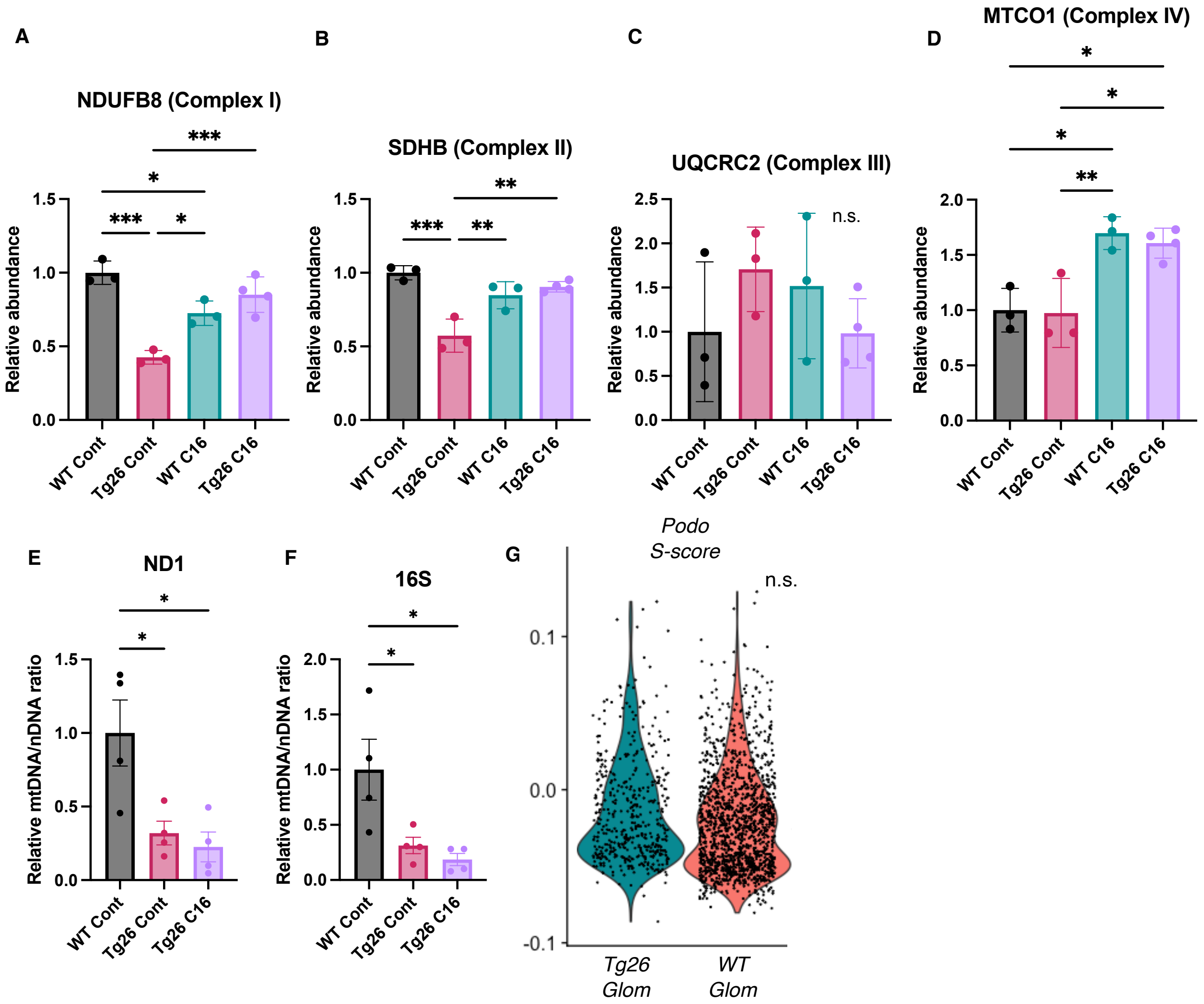


**Supplemental Figure. 4 Additional data about mitochondria and S phase score of podocytes**
(**A-D**) Quantification of Western blot probed against mitochondrial subunits normalized by VDAC.
(**E** and **F**) Relative expression ratio of Nd1 and 16S divided by Hk2. (**G**) Violin plot showing S phase score of Podocytes in each samples.
(One-way ANOVA; *, $P<0.05$; **, $P<0.01$; ***, $P<0.001$)

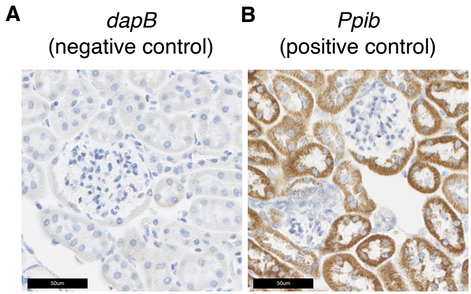


**Supplemental Figure. 5 Additional *in situ* hybridization images**
(**A**) *In situ* hybridization images probing *dapB* (negative control probe) showed no signals.
(**B**) *In situ* hybridization images probing *Ppib* (positive control probe) showed strong signals.

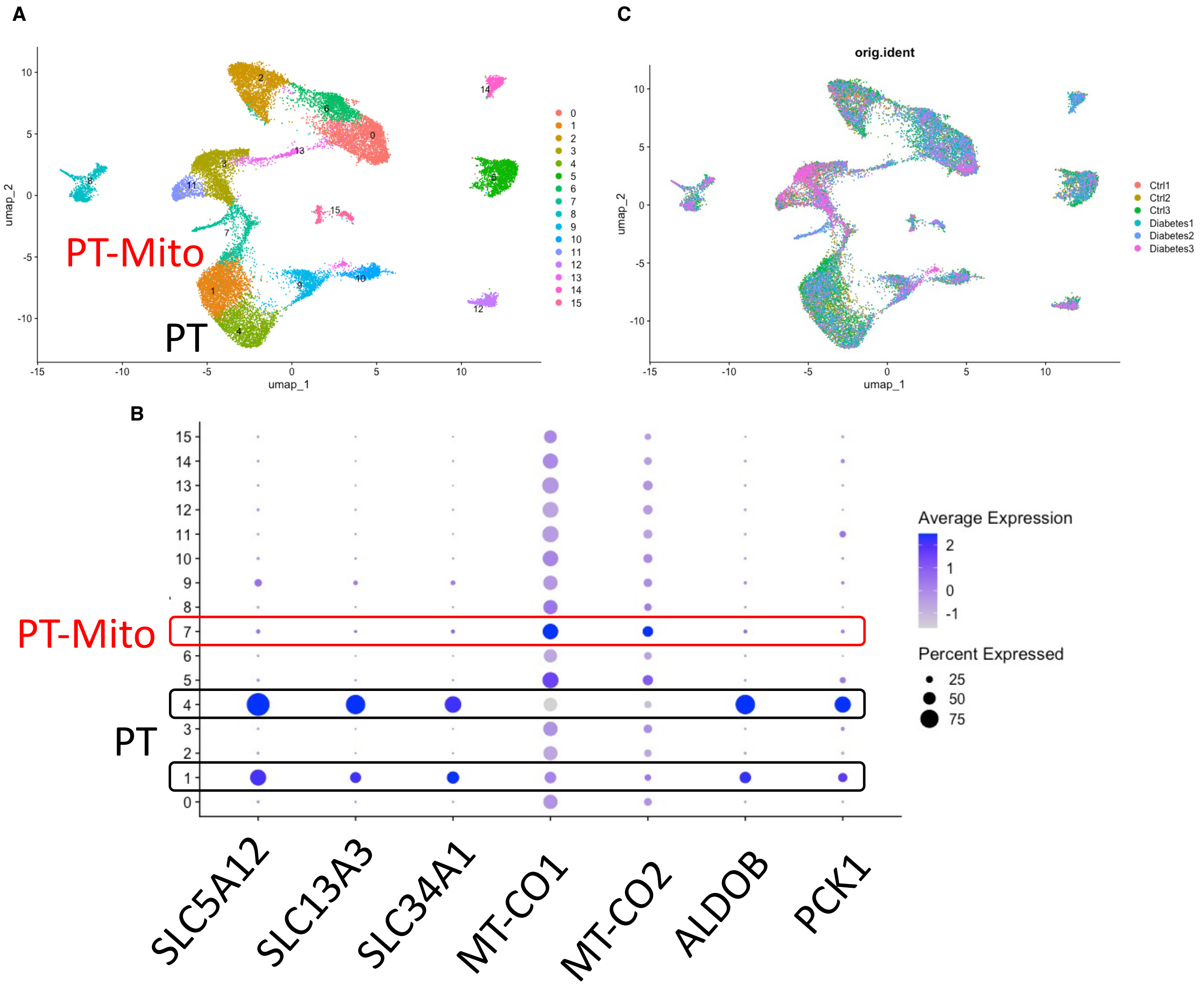


**Supplemental Figure. 6 PT-Mito cluster detection of publicly available human kidney single-nuclear RNA-seq data (GSE131882)**

(**A**) UMAP plot of human kidney single-nuclear RNA-seq data shows 16 clusters. Cluster 1, 4 are proximal tubule (PT) clusters, and cluster 7 is PT-Mito cluster.

(**B**) Dot plot shows expression of PT marker genes and PT-Mito marker genes obtained from current manuscript data. PT-Mito markers including MT-CO1 and MT-CO2 had high expression in cluster 7.

(**C**) UMAP plot shows all six samples are contributing to all cell clusters.

A

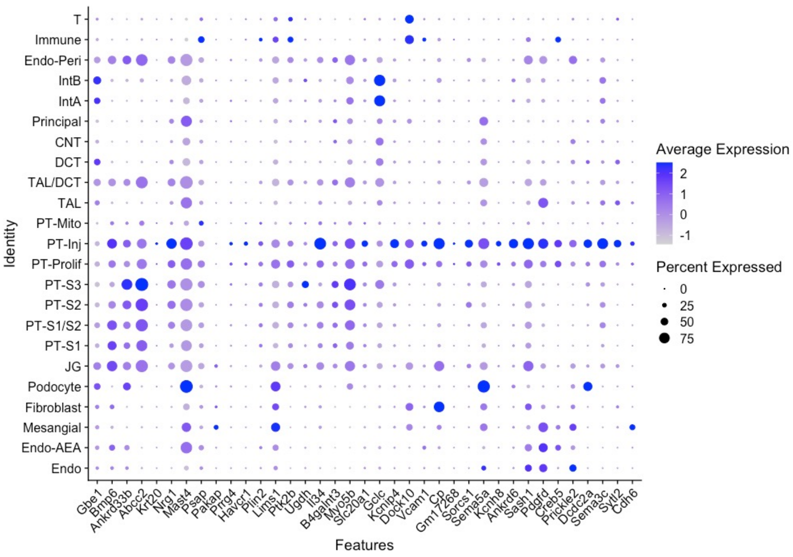


B

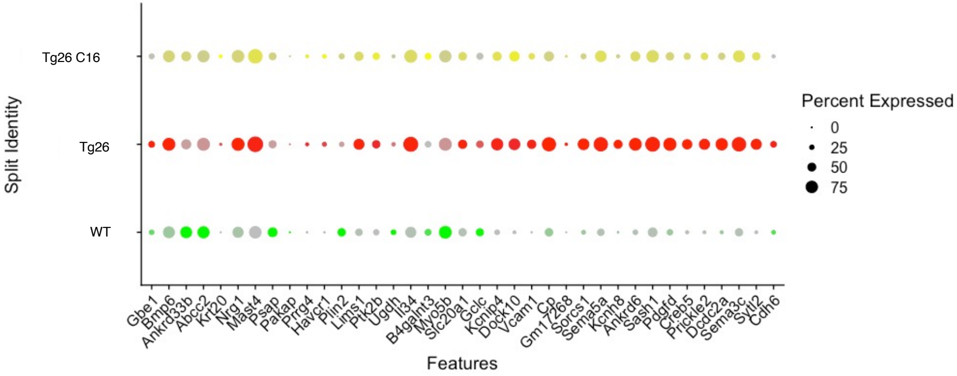


**Supplemental Figure. 7 Single-nuclear RNA-seq data comparison with ischemic reperfusion injury model**
(**A**) Dot plot showing expression of marker genes reported in new proximal tubular clusters by mouse ischemic reperfusion acute kidney injury model (Kirita.PNAS.2020).
(**B**) Dot plot showing expression of marker genes reported (Kirita.PNAS.2020) in PT-inj cells by snRNA-seq

| **Upstream Regulator** | **Activation Z-score WT C16 vs WT** | **Activation Z-score Tg26 vs WT** | **Activation Z-score Tg26 C16 vs Tg26** | **Triple Z-score** |
|---|---|---|---|---|
| **STAT3** | -2.183 | 6.286 | -3.182 | 43.66 |
| **FOXO1** | -3.618 | 3.991 | -2.697 | 38.94 |
| **GLI1** | -2.335 | 5.517 | -2.936 | 37.82 |
| **EPAS1** | -2.665 | 3.554 | -3.242 | 30.71 |
| **SREBF1** | -3.013 | 3.672 | -2.63 | 29.10 |
| **CREB1** | -2.325 | 4.181 | -2.916 | 28.35 |
| **ATF4** | -2.606 | 4.445 | -2.028 | 23.49 |
| **SMAD2** | -2.207 | 2.791 | -2.335 | 14.38 |

**Supplemental Figure. 8 Activation Z-scores from bulk RNA-seq data**
Activation Z-scores of transcription factors by upstream regulator analysis of Ingenuity Pathway Analysis. Triple Z-score was calculated by multiplying three Z-scores in each comparison.

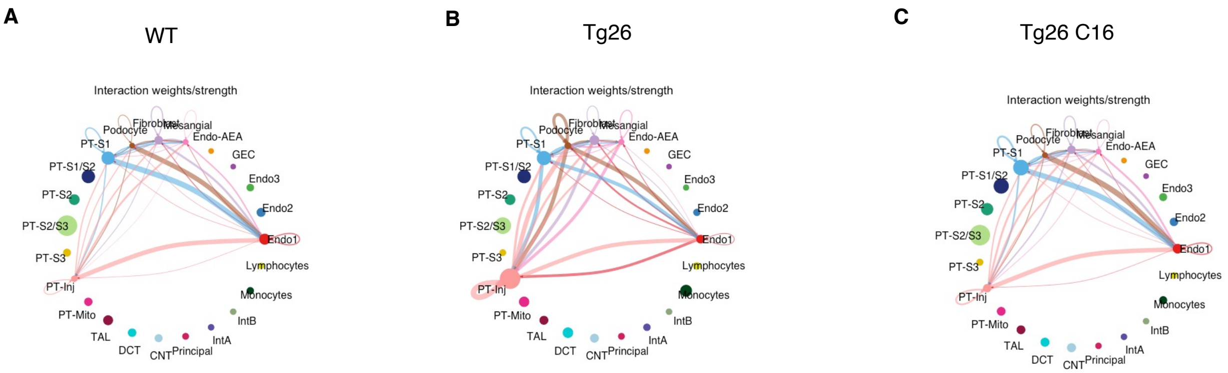


**Supplemental Figure. 9 Additional cell-cell interaction analysis plot**
(**A-C**) Circles plot showing cell-cell interaction weight in each sample at the resolution of each identified cell cluster.
